# Off-purpose activity of industrial and agricultural chemicals against human gut bacteria

**DOI:** 10.1101/2024.09.05.610817

**Authors:** Anna E. Lindell, Stephan Kamrad, Indra Roux, Shagun Krishna, Anne Grießhammer, Tom Smith, Rui Guan, Deniz Rad, Luisa Faria, Sonja Blasche, Nicole C. Kleinstreuer, Lisa Maier, Kiran R. Patil

## Abstract

Contamination by industrial and agricultural chemicals like pesticides are a cause of great concern due to the risk to human and environmental health. While these chemicals are often considered to have restricted activity and are labelled as such, there are concerns over a broader toxicity range. Here we report the impact of 1076 pollutants spanning diverse chemistries and indicated applications on 22 prevalent commensal gut bacteria. Our systematic investigation uncovered 588 interactions involving 168 chemicals, the majority of which were not previously reported to have antibacterial properties. Fungicides and industrial chemicals showed the largest impact with circa 30% exhibiting anti-commensal properties. We find that the sensitivity to chemical pollutants across species surprisingly correlates with that to human-targeted drugs, suggesting common susceptibility mechanisms. Using a genome-wide chemical-genetic screen, we identified membrane transport and fatty acid metabolism as major modulators of the off-target toxicity of chemicals. Mutants exhibiting chemical resistance include those defective in producing human-health-relevant metabolites like branched short-chain fatty acids, indicating that chronic exposure could lead to selection against production of beneficial metabolites. Toxicokinetic modelling suggested gut bacteria could be used as more sensitive *in vitro* toxicity indicators for chemicals of concern than animal models. Together, our data uncovers the off-target activity of industrial and agricultural chemicals with widespread exposure against human gut bacteria. Impact on the structure and function of the microbiota should therefore be considered in assessing chemical safety.

## Introduction

Synthetic chemicals have become indispensable for agriculture and industry. Their pervasive use and the environmental persistence has led to pollution levels that, by some estimates, have already exceeded the safe planetary boundary (Cousins et al. 2022; Steffen et al. 2015; Persson et al. 2022). This presents a critical challenge to both ecosystems and human health. Among the wide range of pollutants, pesticides are a major source of concern, with the FDA and EFSA documenting 212 and 358 different pesticide residues in food respectively (European Food Safety Authority (EFSA), Medina-Pastor, and Triacchini 2020; U.S. Food and Drug Administration 2018), and wider contamination of water sources (Maggi, Tang, and Tubiello 2023; Egli et al. 2023). Other industrial chemicals, such as plasticisers, surface coatings, and dyes, also routinely enter food and water via complex and unintentional exposure routes, either via the environment or food processing and storage (Groh et al. 2021).

The contamination of food and water resources by industrial and agricultural chemicals means that many enter the human gastro-interstinal tract and enterohepatic circulation. Altogether, tens of thousands of xenobiotic compounds are estimated to enter the human body (Lindell, Zimmermann-Kogadeeva, and Patil 2022). E.g., >95% of Americans have detectable levels of per-/polyfluoroalkyl substances (PFAS) in their blood (Calafat et al. 2007). In addition, in a UK cohort all tested urine samples showed detectable levels of pesticides. For example, cypermethrin and permethrin were detected in >96 % of samples, diethylphosphate was detected in 75 % of samples, and glyphosate, a widely used herbicide, was present in 53 % of samples (Smalling et al. 2023). The gut microbiota - a community of hundreds of microbial species - are thus exposed to numerous chemicals on a daily basis. While pharmaceutical drugs have been shown to impact growth and metabolic physiology of gut bacteria (Maier et al. 2018; Weersma, Zhernakova, and Fu 2020), little is known about the impact of agricultural and industrial chemical contaminants (Lindell, Zimmermann-Kogadeeva, and Patil 2022).

Chemicals like pesticides are usually marketed with a narrow definition of target organisms, e.g. as insecticides, herbicides or fungicides. Yet, many studies have shown off-purpose activity on non-target organisms including bacteria. For instance, communities of soil bacteria have been shown to be disrupted by pesticides (Ruuskanen et al. 2023). Several pesticides have been reported to exhibit potential anti-pathogenic activity (Chen et al. 2018; Tran et al. 2016; Lim et al. 2013; Clarke et al. 2023). A small number of pesticides and industrial chemicals have also been shown to impact human and animal gut microbiota (Giambò et al. 2021; Utembe and Kamng’ona 2021; Gois et al. 2023; Mesnage et al. 2022; Hu et al. 2021; Mesnage et al. 2021; Ali and AlHussaini 2024; Defois et al. 2018; Riesbeck et al. 2022). While these previous studies indicate that many pesticides and other chemical pollutants might exhibit off-purpose activity against commensal gut microbiota, there is a notable lack of systematic assessment. Toxicological safety assessments for these chemicals do not consider the gut microbiome. Assessing this *in vivo* is difficult since controlled studies cannot be done from an ethical standpoint and observational studies are hampered by multiple confounding factors in population studies.

We therefore embarked on a large-scale screen assessing the anti-commensal effect of 1076 compounds consisting of pesticides, pesticide metabolites, and industrial chemicals. This bottom-up approach allows creating a knowledge base and mechanistic insights that can be used for hazard identification and to design observational studies. Indeed, establishing drug usage as one of the top variables underlying inter-individual microbiome variation was accelerated by the bottom-up interaction map between therapeutics and gut bacteria (Maier et al. 2018; Klünemann et al. 2021). Using a comprehensive chemical library and a panel of representative human gut bacteria, we uncover numerous growth inhibitory interactions, particularly with fungicides and industrial chemicals, and kinetic modelling reveals significant effects at environmentally relevant exposures. The identified interaction network and the follow-up genetic analysis substantially expands the knowledgebase on xenobiotic toxicity in bacteria, revealing common principles and paving the way for computational predictions.

## Results

### Comprehensive library of chemical contaminants

To systematically assess the impact of chemical contaminants, we used an extensive library of 1076 compounds, many of which are likely to enter food and water due to pollution, agricultural application or industrial processing (**Figure 1A+B**). The library contains 829 pesticides, mainly herbicides, insecticides and fungicides. Beyond these widely used major classes, it contains a diverse set of pesticides with other target organisms including spiders, nematodes, bacteria and rodents. Furthermore, the library contains 48 industrial chemicals. These include polycyclic aromatic hydrocarbons (PAHs) which can be introduced into foods during cooking at high temperatures, bisphenol compounds that can leach into food from packaging materials and nitrosamines which have toxic and carcinogenic potential and are a common contaminant in medications. Several chemicals in the library are classified as persistent organic pollutants, e.g. PFAS as well as the insecticides DDT and chlordecone. The library also includes 119 known pesticide metabolites, which are important to consider for their toxicological effects as they can arise from biotransformation either outside or inside the human body (Koppel, Maini Rekdal, and Balskus 2017). The library further included 75 pesticide-related compounds, such as synthetic precursors or breakdown products, pesticide isomers not themselves used as pesticides, or products that might be included in pesticide formulations. Finally, the library includes 5 mycotoxins which might be found in mould-contaminated foods. Our library covers the vast majority (291 out of 352) of pesticide classes described in the British Crop Production Council (BCPC) Pesticide Compendium and 87 % of compounds described in a recent paper highlighting pesticide movement along rivers into oceans (Maggi, Tang, and Tubiello 2023). This library, together with extensive metadata we compiled (**Supplementary Data 1**), constitutes a chemically diverse resource for systematic investigation of the impact of chemicals of concern on biological systems.

**Figure 1:**
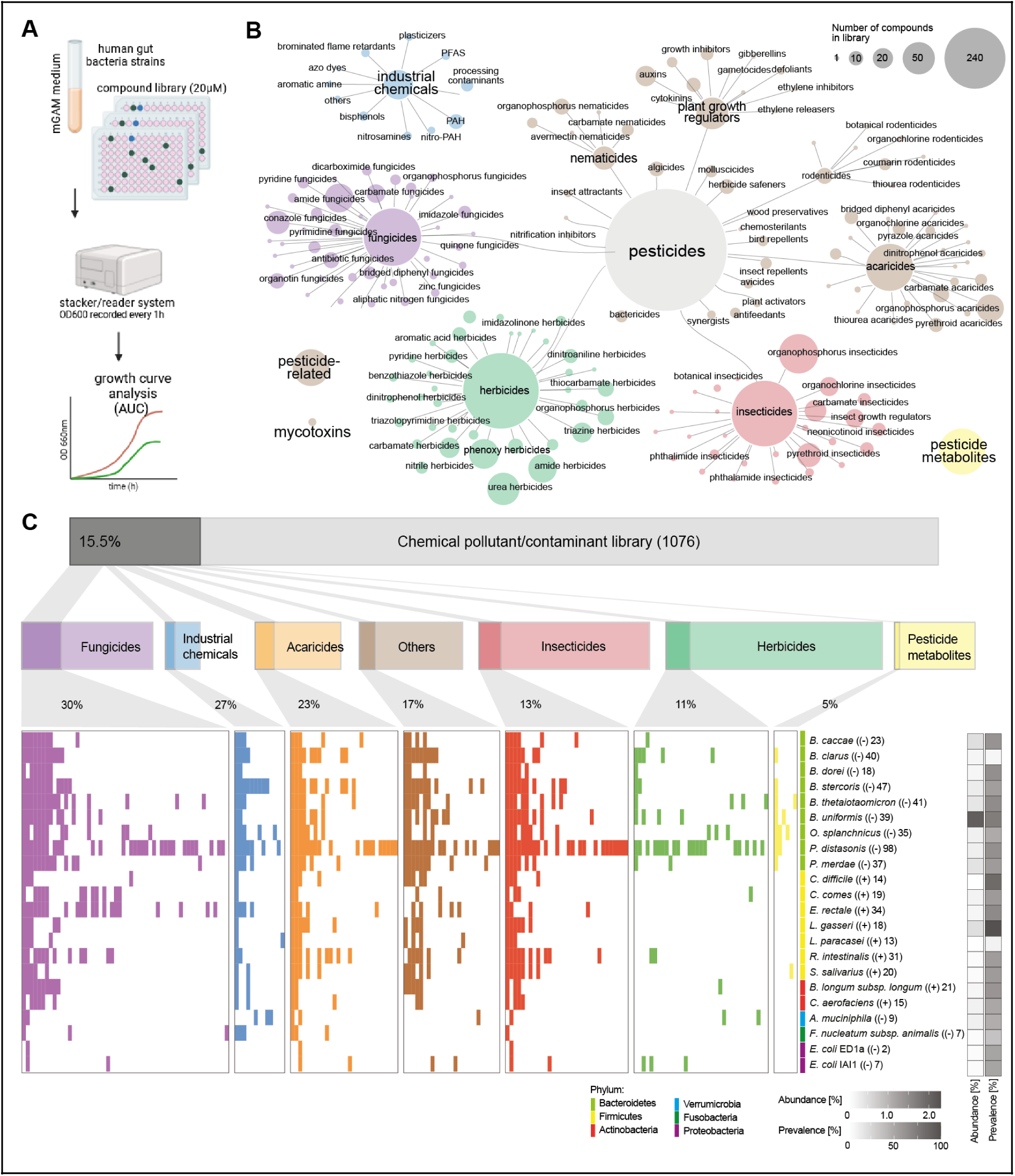
Extensive impact of food contaminants on human gut commensals. (A) High-throughput screening workflow to test the susceptibility of gut bacteria against industrial and agricultural chemicals. Mono-cultures of commensal gut bacteria were grown in mGAM medium with 20 μM of each compound (n=3 biological replicates). Growth curves were recorded and the area under the curve (AUC) was used as a fitness proxy. (B) A library of xenobiotic agricultural and industrial chemicals was assembled in 96 well plates. The network illustrates the number of library compounds targeted at different organisms and their chemical subclasses, based on the BCPC Pesticide Compendium. Out of 1076 compounds in total, 829 were pesticides, 119 were pesticide metabolites, 48 were industrial chemicals, 5 were mycotoxins and 76 were other, pesticide-related compounds such as synthetic precursors or compounds that might be found in commercial formulations. (C) High-throughput growth screening was used to assess the impact of food contaminants on individual gut bacterial strains. Significant pesticide-strain interactions, where the growth of a specific strain is inhibited versus DMSO control, are highlighted with a coloured bar (p_adj_<0.05, >20% reduction in AUC, 2 of n=3 biological replicates significant). Phylum, abundance and prevalence of strains is displayed through colour and grey-scale bars, and gram-stain and number of compounds that inhibit growth are added in parenthesis after strain name.

### Chemical contaminants from diverse classes inhibit growth of commensal gut bacteria

For all 1076 compounds in the library, we assessed their impact on the growth of 22 gut bacterial strains belonging to 21 species (**Figure 1C**). These strains were selected based on their abundance and prevalence in the healthy human gut microbiota, and phylogenetic and metabolic representation (**Supplementary Data 2**) (Tramontano et al. 2018; Dai et al. 2022). All strains were grown anaerobically at 37°C in mGAM broth, a rich medium designed to support the growth of a maximum number of strains. All screens were performed in 96 well plates and at a compound concentration of 20 μM as described previously (Müller et al. 2024). A concentration of 20 μM was chosen to allow direct comparison with previous studies (Maier et al. 2018). Bacterial growth was monitored for 24 h and quantified as the area under the growth curve (AUC). FDR-corrected z-score statistic (p_adj_<0.05) and a fold-change cut-off of >20% was used to identify compounds which significantly and substantially decreased the growth of the tested bacterial strains in at least two out of three biological replicates (**Supplementary Data 3**, **Supplementary Figure 1A**).

Out of the 1076 chemicals tested, 168 (15.5 %) were active against at least one bacterial strain (**Supplementary Figure 1B**). From the strain perspective, we observed varied sensitivity; Bacteroidetes, especially *Parabacteroides distasonis*, were the most sensitive while *Escherichia coli*, *Fusobacterium nucleatum subsp. animalis* and *Akkermansia muciniphila* were the least sensitive (**Supplementary Figure 1C**). Chemical susceptibility profiles of bacterial species grouped clearly by evolutionary relatedness, i.e. by phylum (**Supplementary Figure 2B**). Accordingly, most anti-commensal compounds showed clear differences in toxicity in different phyla (**Figure 2A**). E.g., some Firmicutes were uniquely sensitive to a group of conazole fungicides, including the widely used compounds imazalil and prochloraz. This indicates a shared genetic basis for toxicity and suggests that chemical susceptibility data could be used to predict effects on related bacterial species for *in silico* toxicological assessments.

**Figure 2:**
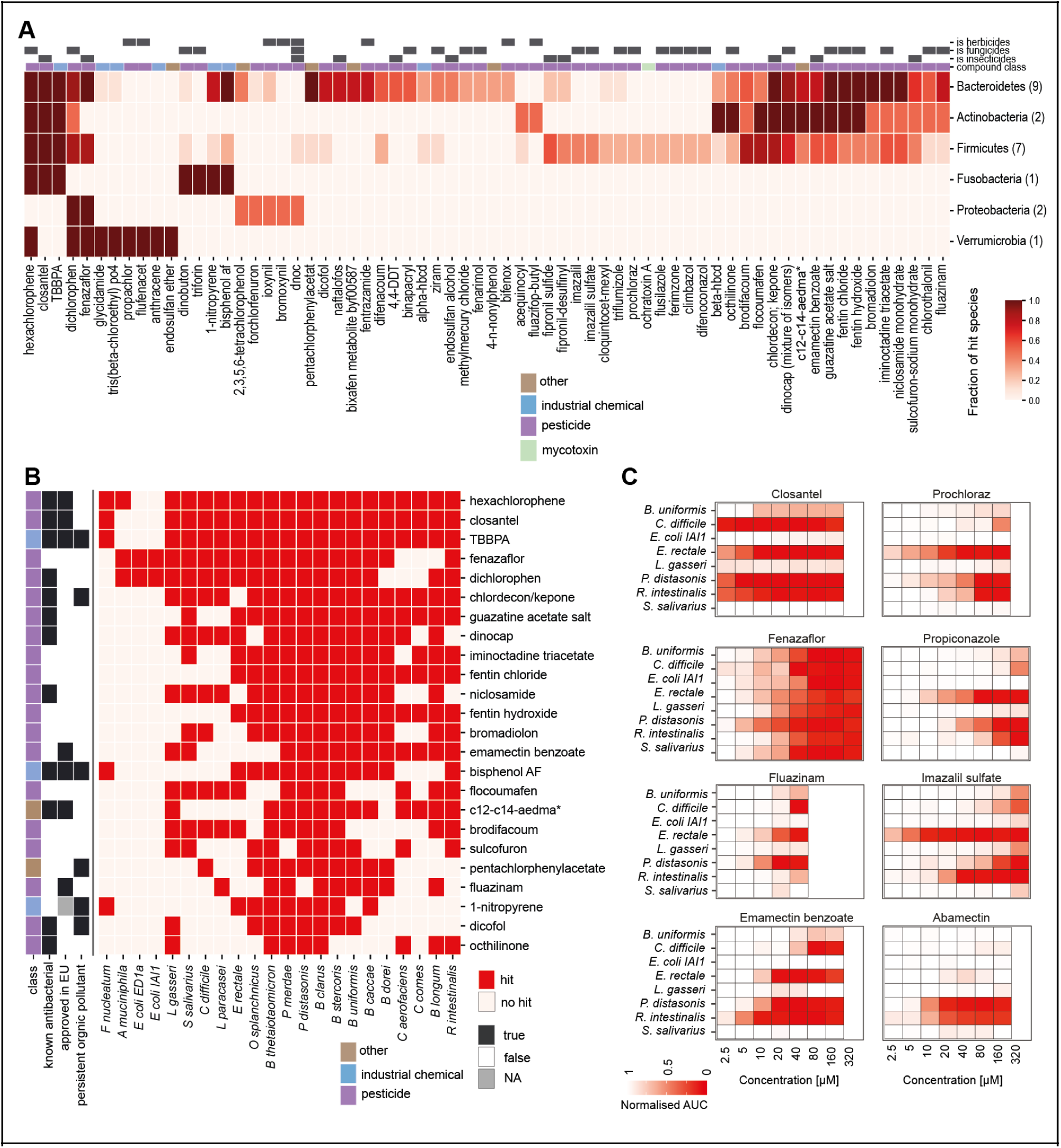
Screen identifies broad-spectrum anti-commensals. A) Chemical susceptibility profiles grouped by phylum. Compounds (x-axis) and phyla (y-axis) were clustered by profile similarity, using Euclidean distance and Ward clustering. The number of strains in each phylum is indicated in parentheses. B) Heatmap showing growth inhibition by compounds (20 μM) which were identified to have broad-spectrum activity, inhibiting more than a third of the tested species. Results shown are a subset of those shown in Figure 1C. Several of these compounds are widely used and/or persistent organic pollutants, raising concerns about their potential anti-commensal effects. Abbreviation: *C_12_-C_14_-Alkyl(ethylbenzyl)dimethylammonium chloride. C) Result of independent validation screen. Selected bacteria and compounds were screened for growth interactions at 8 different concentrations (n=3 biological replicates). Colours indicate the relative AUC compared to DMSO controls.

On the chemical side, fungicides, industrial chemicals and acaricides had >20 % prevalence of compounds showing growth inhibitory effects (**Figure 1C**). Most compounds were effective against a few strains indicating a specific, narrow spectrum impact. Yet, 24 compounds showed a broad toxicity range inhibiting more than a third of the species tested (**Figure 2B**). Some of these compounds have known antimicrobial properties, such as hexachlorophene which is used to disinfect skin prior to surgery. Three 4-hydroxycoumarin-like vitamin K antagonist rodenticides of the warfarin family were also highly toxic to most bacteria. Amongst the industrial chemicals, two structurally related bisphenols, tetrabromobisphenol A (TBBPA), a brominated flame retardant, and bisphenol AF, used in plastics, exhibited broad-spectrum toxicity. Among pesticides showing broad spectrum activity, many are no longer approved in the EU, but many appear to be still in use in the US, namely niclosamide, fentin hydroxide, bromadiolon, brodifacoum and octhilinone. Among those approved in the EU were emamectin benzoate, a widely used macrocyclic lactone insecticide, and fluazinam, a widely used fungicide. The toxicity of emamectin benzoate is likely due to the presence of benzoate, a common antibacterial preservative, as the closely related pesticide abamectin (which was not formulated as a benzoic acid salt) only inhibited two species. Several of the broad-spectrum anti-commensal compounds are classified as persistent organic pollutants and are therefore likely to enter the human food chain via the environment, even when no longer actively in use. These included the insecticides dicofol, pentachlorophenyl acetate and chlordecon, the plasticiser bisphenol AF (BPAF) and the flame retardant TBBPA. Overall, these results indicate that several widely used and environmentally persistent industrial and agricultural chemicals show broad anti-commensal activity.

To assess the potency of the anti-commensal chemicals, we determined minimum inhibitory concentrations (MIC) for 8 pesticides and 8 bacteria (**Figure 2C**). Independently purchased compounds were used to account for any errors associated with batch variations and large-scale library preparations. The majority of the interactions from the main screen (75%, 24 out of 32) were also observed in this independent experiment (**Supplementary Figure 1D+E**). Many compounds show strong inhibitory effects at substantially lower concentrations than were tested in the main screen. E.g., imazalil sulfate and prochloraz inhibited *Eubacterium rectale*, a prevalent species that produces health-associated metabolite butyrate, at a concentration as low as 2.5 µM. Taken together, our systematic investigation revealed hundreds of new chemical-bacteria interactions including chemical contaminants with broad and potent anti-commensal activity.

### Chemicals alter composition of synthetic gut bacterial communities

We next investigated how species-level effects of chemicals translate in bacterial communities using a synthetic community of 20 gut bacteria (Com20, Methods, (Müller et al. 2024)) (**Figure 3A, Supplementary Data 4**). Communities with an equal initial ratio of all species were cultivated and transferred twice in fresh growth medium (mGAM) before being challenged with bisphenol AF (BPAF) or TBBPA - two chemicals with broad-spectrum anti-commensal activity. For BPAF, we observed compositional changes concordant with the effects observed in mono-culture. These include decreased relative abundance of *Bacteroides thetaiotaomicron, Bacteroides uniformis*, *Parabacteroides merdae*, and *Roseburia intestinalis*. Species that were insensitive in mono-culture showed an increased abundance, viz. *Collinsella aerofaciens*, *Coprococcus comes* and *Streptococcus salivarius*. Two species, *F. nucleatum* and *Eubacterium rectale* were sensitive in mono-culture but were protected in the community (**Figure 3B**). In the case of TBBPA, community structure was characterised by the monodominance of *B. thetaiotaomicron*, despite this species being susceptible in mono-culture. These experiments indicate that xenobiotics which inhibit single strains in mono-culture also inhibit communities with individual species’ sensitivities reflected in relative abundance changes in communities to certain degrees as well as community-scale effects like cross-protection. These observations are concordant with the effects observed in gut bacterial communities exposed to therapeutic drugs (Garcia-Santamarina et al. 2023; Klünemann et al. 2021). In the case of therapeutic drugs, the emergent community-scale effects are due to bacterial metabolism of drugs and the impact of drugs on bacterial metabolism. The results from the BPAF and TBBPA suggest that similar mechanisms are applicable in the case of chemical contaminants.

**Figure 3:**
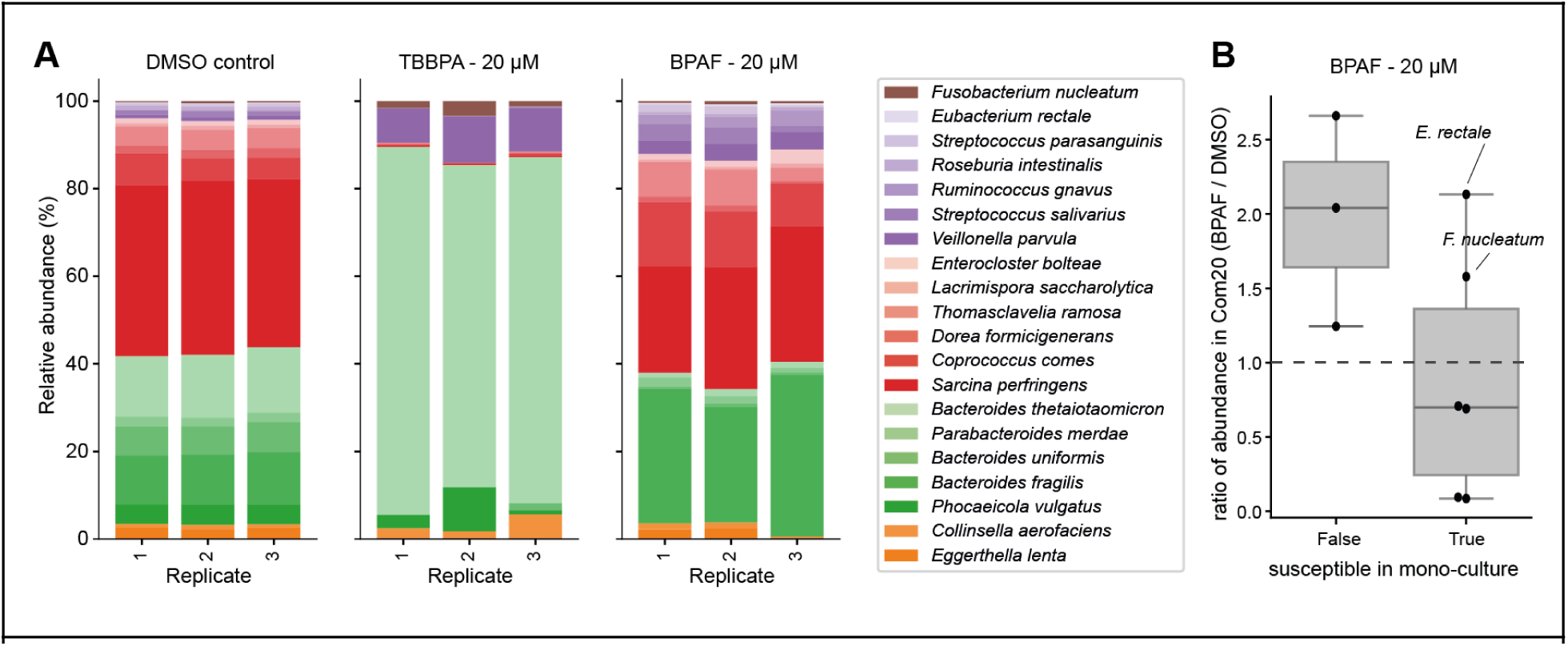
Impact of bisphenol pollutants on community structure. (A) A synthetic community of 20 bacterial species (Com20) was grown *in vitro* in the presence of bisphenol AF (BPAF) and tetrabromobisphenol A (TBBPA) and community composition was assessed by 16S amplicon sequencing. Relative abundances of species, coloured by phylum, are shown. The two compounds, which showed strong effects on mono-cultures, also showed strong effects in the community. (B) Comparison of mono-culture versus community susceptibility to BPAF. The absolute abundance of each species in the community was determined from 16S amplicon sequencing counts and overall culture optical densities. Shown is the ratio of abundances between cultures treated with BPAF versus DMSO control. Both approaches returned largely similar results, although *E. rectale* and *F. nucleatum* stood out as they were protected in the community but susceptible in mono-culture.

### Toxicokinetic modelling shows that gut bacteria could be used as sensitive indicators of chemical toxicity

We applied physiologically based toxicokinetic (PBTK) modelling to a subset of 168 library compounds for *in vitro* to *in vivo* extrapolation (IVIVE). Using a 30 day exposure scenario with 24 hour dose intervals, we estimated equivalent administered oral doses in humans (EADs) that result in concentrations of xenobiotics in the gut at which we observed bacterial growth inhibition *in vitro* (**Supplementary Data 5**). We compared these EADs to the doses at which toxicity was observed in a curated database of animal experiments (ToxValDB). This comparison is captured as the POD_ratio_, which is the ratio of lowest dose at which an adverse effect was observed in mice (Point of departure - lowest adverse effect level: POD-LOAEL) to the previously determined EAD. We detected a positive log_10_(POD_ratio_) for the vast majority of compounds in our analysis (56 out of 68 compounds for which animal data was available) (**Supplementary Figure 3**). A positive log_10_(POD_ratio_) indicates that our *in vitro* screen picked up effects at lower equivalent doses than required to elicit toxicity in animals. While this is not unexpected, as animal toxicology does not usually consider the microbiome, this indicates that *in vitro* screening can be a sensitive method to assess potential adverse effects on microbiota not typically picked up in animal toxicity studies. More consideration and innovative study designs are required to better capture the microbiota-damaging toxic effects of xenobiotics in animal studies.

### Expanded coverage of xenobiotic interaction landscape

Large-scale surveys of xenobiotic-bacteria interactions and computational analyses could enable *in silico* toxicity predictions for xenobiotics (McCoubrey et al. 2021), if the chemical space is sufficiently covered. We therefore compared the Extended-Connectivity Fingerprints (ECFP) of our pollutant library against a random selection of 250,000 compounds from PubChem and a previously assessed library of 1,400 pharmaceutical drugs (Maier et al. 2018) (**Figure 4A**). We note that chemical pollutants are chemically distinct from the set of previously investigated pharmaceutical drugs. Excluding 30 compounds which were part of both libraries, only 6 pollutants (0.6%) had a Tanimoto similarity (a metric which measures the overlap of molecular fingerprints (Maggiora et al. 2014)) > 0.75 to a previously investigated drug (**Figure 4B**). We also compared the Tanimoto similarity index (Bajusz, Rácz, and Héberger 2015) between the PubChem compounds and the screened compound in the libraries and observed that the index for the closest compound was significantly increased in the combined library (p < 2.2e-16; KS test, **Figure 4C**), indicating a greater overall coverage of the chemical space, which could be used to predict the impact of novel compounds.

**Figure 4:**
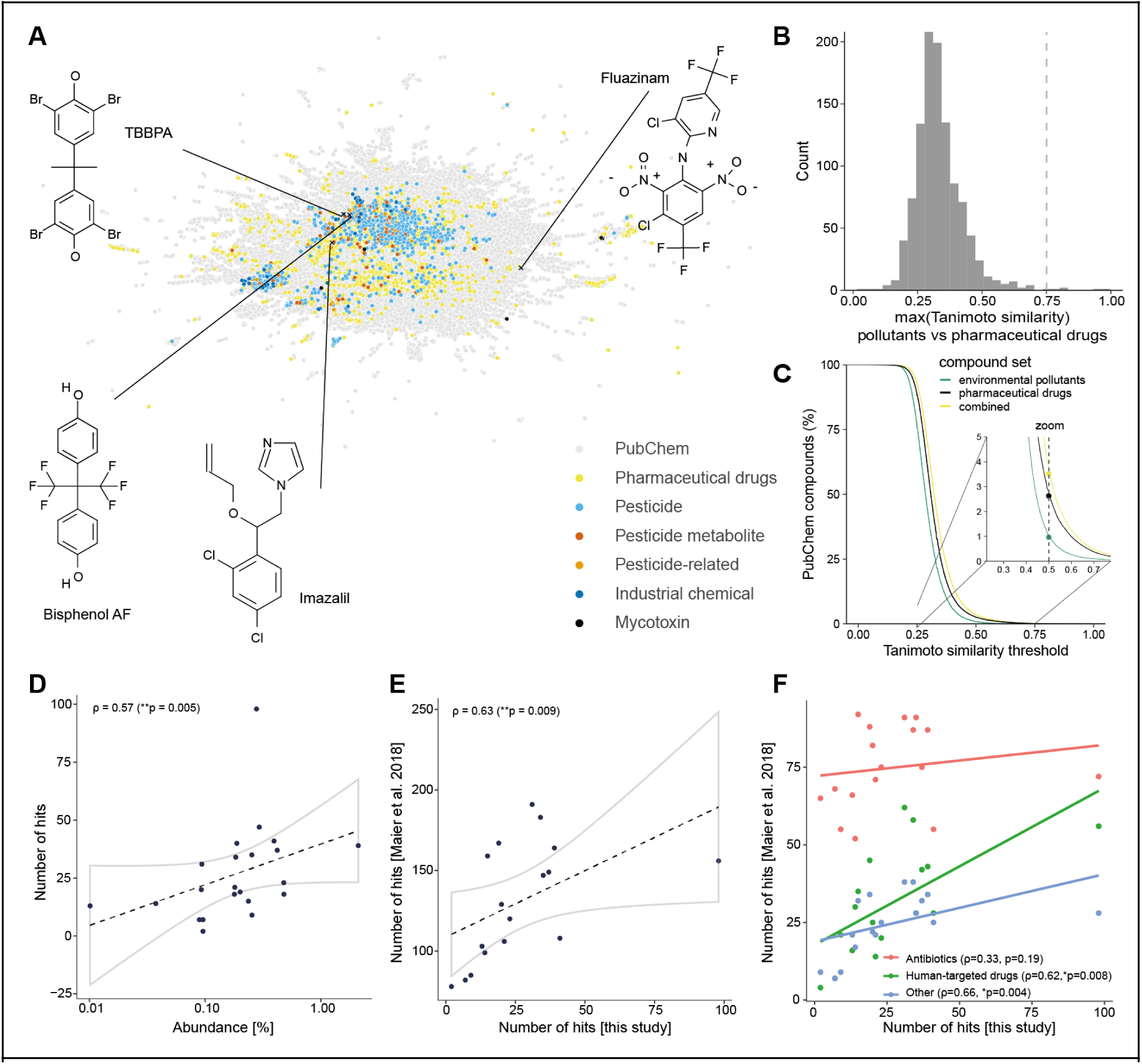
The xenobiotic-bacteria growth interaction landscape. A) Uniform Manifold Approximation and Projection (UMAP) for Extended-Connectivity Fingerprints (ECFPs) of library compounds from this study, previously screened pharmaceutical drugs (Maier et al. 2018) and a random selection of 250,000 compounds from PubChem. B) Distribution of similarities between closest matching pollutants from this study and drugs used by Maier et al. 2018. The two compound sets are structurally diverse and distinct, as only very few (6) compounds from the library used in this study had a similarity >0.75 to a drug from (Maier et al. 2018). Compounds shared between the two studies (n=32, mostly veterinary drugs) were excluded from this analysis. C) Coverage of chemical space is increased by our new, complementary xenobiotic compound library. D) Correlation between average species abundance in human microbiome and compound sensitivity. For each strain the abundance is plotted against the number of compounds that inhibited the growth of that strain. Environmental pollutants inhibit the growth of more abundant species more than that of less abundant species. Lines depict the best linear fit including the confidence interval. The Spearman rank correlation coefficient (ρ) and corresponding p-value are displayed. E) Correlation between the number of inhibiting compounds for a specific strain in the Maier et al. 2018 drug screen and this study. Strains that were more sensitive in the Maier et al. 2018 drug screen were also more sensitive to the tested pollutants. F) Correlation between the number of inhibiting compounds for a specific strain in the Maier et al. 2018 drug screen and this study grouped by drug type. Sensitivities to human-targeted and ‘other drugs’ from Maier et al 2018 correlate significantly with sensitivity to environmental pollutants tested in this manuscript. Sensitivity to antibiotics in Maier et al 2018 is not significantly correlated with sensitivity to environmental pollutants.

### Common factors likely underpin susceptibility and resistance to therapeutic drugs and chemical contaminants

We noted a positive correlation of number of compounds affecting a species with its abundance in the human gut (Spearman rank correlation: ⍴ = 0.57, p-value = 0.005) (**Figure 4D**), but none with the prevalence (**Supplementary Figure 2A**). Thus, chemical pollutants impact abundant species more than low abundant ones suggesting that compounds with broad as well as limited spectrum activity on gut bacteria can cause ecological instability in the microbiota, the former through affecting many species, the latter through affecting few but abundant species with high contribution to the structure and function of the community The observed correlation between abundance and sensitivity to chemicals bears a striking resemblance to that observed previously for therapeutic drugs (Maier et al. 2018, 2021). The strains identified in the drug screen as being sensitive to many compounds were also sensitive to many chemical pollutants tested in this study (⍴ = 0.63, p-value = 0.009) (**Figure 4E**). When stratifying the drugs as antibiotics and human-targeted drugs, the correlation with antibiotics was weaker (⍴ = 0.33, p-value = 0.19) than that with the human-targeted drugs (⍴ = 0.66, p-value = 0.004) (**Figure 4F**). This is consistent with the antibiotics generally exhibiting a broader spectrum activity against commensals than both drugs and chemical pollutants. The correlation in the case of human-targeted drugs indicates that bacteria use a common and non-specific mechanism to tackle non-antibiotic drugs and chemical pollutants.

### Forward genetic screen uncovers chemical selection for loss of beneficial metabolite pathways

To identify the genes modulating the impact of xenobiotics on bacterial fitness, we used a pooled transposon mutant library of *P. merdae* to conduct a competition assay against a panel of 10 chemicals (Voogdt et al. 2024). The library encompasses barcoded transposon mutants of >3000 non-essential genes enabling a system-wide analysis. The library was grown until the early stationary phase and the selection was quantified by using TnBarSeq (Barcoded Transposon Sequencing) (**Supplementary Data 6**). The compounds were selected based on exhibited inhibition of *P. merdae* at concentration of 20 μM or less: TBBPA, BPAF, imazalil sulphate, fluazinam, closantel and emamectin benzoate. For the remaining three compounds (glyphosate, propiconazole, PFOA, PFNA), a larger concentration was tested to impose growth fitness selection. We defined TnBarSeq gene hits as an absolute log_2_(fold-change) > 0.25 and padj < 0.05 compared to the end-point control (DMSO or media). TBBPA, BPAF and Closantel exhibited the largest number of gene hits for compounds tested at ≤ 20 μM, while PFNA 500 μM showed the largest total number of hits (**Supplementary Figure 4A**).

Closantel at 2.5 μM had the biggest impact in library growth and genetic selection, and outgrowth was composed of > 90 % of mutants corresponding to > 20 different insertion positions in the locus corresponding to a single gene, NQ542_01170 (**Figure 5A**). NQ542_01170 encodes a transcriptional regulator homologous to the efflux repressor *acrR* from *B. uniformis*, which is also genomically co-located with *tolC* and RND efflux genes (Hibberd et al. 2017). In *B. uniformis acrR* mutants present increased efflux and resistance to chemical stress, and a similar efflux reconfiguration might drive resistance to closantel in *P. merdae*. TBBPA at 20 μM also shows a strong positive selection for *acrR* mutants, with over a 100-fold increase compared to DMSO control (**Figure 5A**). Interestingly, *acrR* is not a hit for the rest of the pollutants examined (**Figure 5B**). This suggests a degree of specificity in the efflux of chemicals and/or that efflux alone is insufficient in tackling the other chemicals.

**Figure 5:**
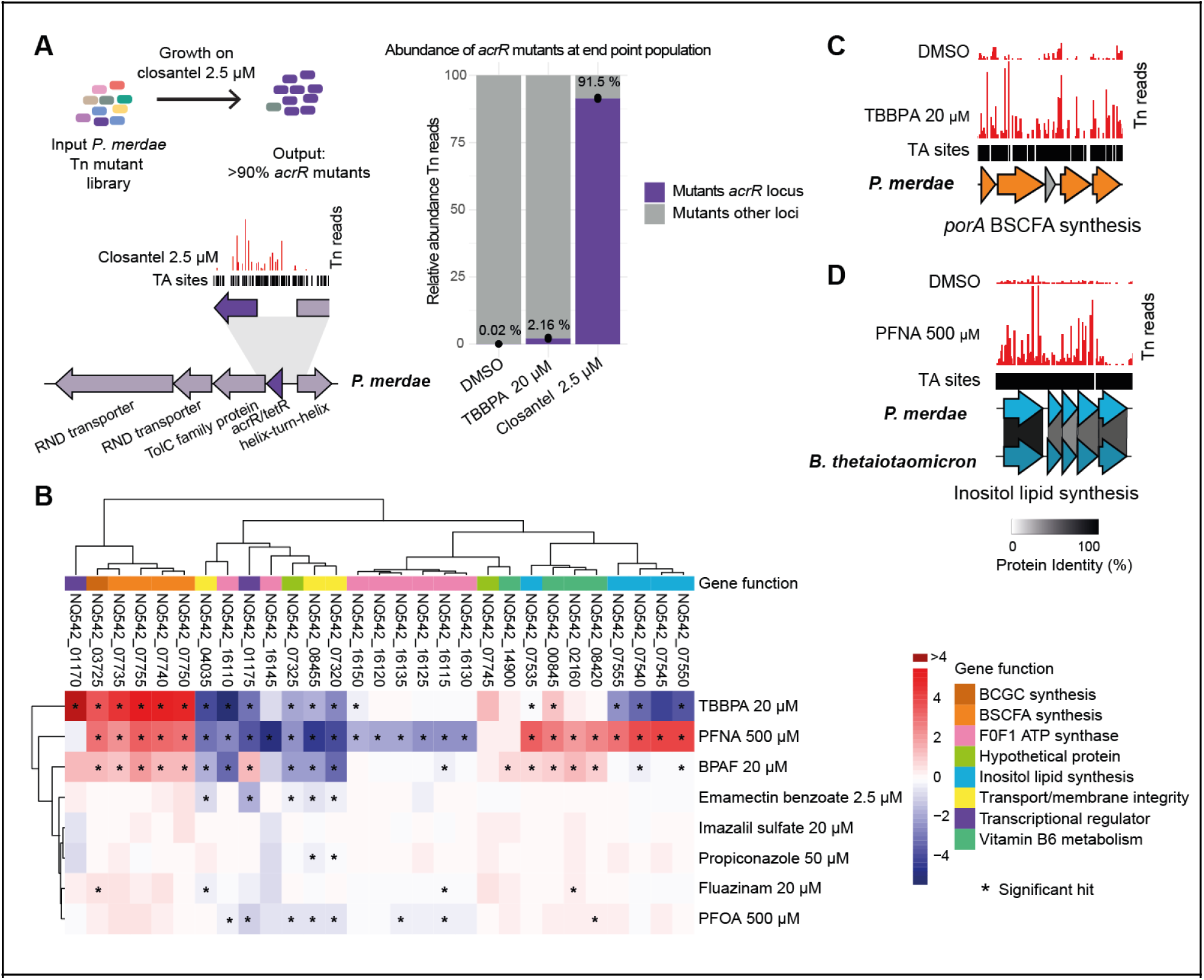
Transposon chemical-genetic screen in *Parabacteroides merdae* reveals genes that modulate susceptibility and resistance to xenobiotics. A) Relative abundance of transposon mutants on the putative *acrR* efflux regulator locus shows strong positive selection under closantel 2.5 μM and TBBPA 20 μM. Representative TnBarSeq insertion plot for closantel and overview of *acrR* genetic context. Coloured arrows represent gene direction and length. Black track above the gene represents TA sequence sites, which are potential Mariner Transposon insertions sites. Red lines represent TnBarSeq insertion sites and read count. B) Heatmap of representative top gene hits. The colour represents the log_2_(fold-change) in Tn mutants with red indicating positive conditional fitness and blue conditional negative fitness compared to the end point DMSO control. Significant hits padj<0.05 abs log_2_(fold-change)>0.25 are shown with an asterisk. Gene function categories are indicated in colours. C) Representative TnBarSeq insertion plot for the *porA* branched short-chained fatty acid gene cluster (orange) in TBBPA 20 μM and DMSO control. D) Representative TnBarSeq insertion plot on the inositol lipid biosynthesis gene cluster in PFNA 500 μM and DMSO control. Homology to the characterised inositol lipid gene cluster from *B. thetaiotaomicron* is shown using Clinker.

The next top hits for positive selection under TBBPA 20 μM were four genes corresponding to the *porA* catabolic gene cluster with a >30-fold enrichment. This pathway is involved in the degradation of branched chain amino acids (BCAAs) into branched short-chain fatty acids (BSCFAs) (Qiao et al. 2022) (**Figure 5C**). In Qiao et al, the *P. merdae porA* mutant (NQ542_07740, pyruvate:ferredoxin oxidoreductase) lost its beneficial probiotic property of preventing atherosclerosis in a mouse model. PorA is present in several gut bacteria and conserved across Gram + and Gram - bacteria, including *Clostridium sporogenes* where it is also involved in the catabolism of other amino acids and plays a role in immunomodulation (Dodd et al. 2017; Guo et al. 2019). Furthermore, KEGG pathway analysis over all gene hits under TBBPA selection showed a significant enrichment in BCAA degradation genes (**Supplementary Figure 4B**), including a putative BCAA aminotransferase (NQ542_03725). This gene is a close homolog of *B. fragilis* BF9343–3671 (74% identity, 100% coverage) whose knock out mutant stops producing immunomodulatory branched-chain α-galactosylceramides lipids (GCs) impacting colonic NKT cell regulation in mono colonised mice (Oh et al. 2021). Besides TBBPA, NQ542_03725 transposon mutants are also a hit for positive fitness enrichment in PFNA 500 μM, BPAF 20 μM and Fluazinam 20 μM and *porA* gene cluster is also a positive selected hit for PFNA 500 μM and BPAF 20 μM (**Figure 5B**). If these xenobiotics select for the loss of bacterial biosynthesis of branched amino acid-derived beneficial metabolites such as BSCFAs and GCs *in vivo,* they could further impact human health via gut-heart or gut-immune axes.

Loss-of-function mutants of the accessory metabolism genes NQ542_07535–55 were positively selected under PFNA 500 μM challenge (circa 16-fold enrichment). These genes were identified as a putative inositol and inositol lipid biosynthetic gene cluster by homology to *B. thetaiotaomicron* (BT_1522–6) (**Figure 5D**). This inositol lipid gene cluster is widespread in Bacteroides, and loss of inositol lipid production in *B. thetaiotaomicron* was associated with changes in cell capsule and increased resistance to antimicrobial peptides (Heaver et al. 2022). Interestingly, we also observe that these genes are among the main negative fitness hits under TBBPA 20 μM (**Figure 5B**). Finally, BPAF 20 μM presented a significant enrichment in hits mapping to the KEGG pathway for vitamin B6 metabolism, with positive selection hits also under PFNA 500 μM (**Figure 5B**). In summary, TnBarSeq shows modulation of metabolism via disruption of accessory metabolism pathways can confer increased resistance to some chemical pollutants.

### Membrane and transport pathways modulate xenobiotic resistance

Among the top sensitive genes across xenobiotics are the membrane proteins NQ542_07320 and NQ542_08455 (negative fitness in 7 out of 10 xenobiotics) from the FadL family of hydrophobic compound transporters (**Figure 5B**). When analysing the subset of genes encoding transporters, the RND multidrug efflux locus NQ542_073240–45 coding a putative AcrAB/TolC also stands as a common gene with negative fitness (**Supplementary Figure 4C**). Another common hit for low fitness are the genes from F0F1 ATP synthase complex, whose loss-of-function has been associated with increased sensitivity to antimicrobial peptides in anaerobic conditions (Vestergaard et al. 2017). This is also consistent with bisphenol exposure leading to overexpression of F0F1 ATP synthase subunits in *B. thetaiotaomicron* (Riesbeck et al. 2022). Beyond transporters, the putative lipid A phosphoethanolamine transferase NQ542_04035 of the *eptA* gene family is important for fitness (5/10 conditions). This class of enzymes often mediates resistance to positively charged host antimicrobial peptides and polymyxin antibiotics via modifications in membrane polarity (Samantha and Vrielink 2020). Other relevant genes for xenobiotic survival are membrane protein assembly components and chaperones (Skp and Bam family proteins) (**Supplementary Figure 4C**). Overall, genes implicated in efflux, membrane polarity and membrane integrity are important for survival across diverse xenobiotics.

## Discussion

The gut microbiota are a neglected potential site of action for ingested xenobiotics due to presumed specificity of these compounds. For example, fungicides with fungi-specific targets are not expected to impact bacteria. Yet, the vast genetic diversity of the gut microbiome means that the microbiome may harbour vastly more potential xenobiotic targets than even the human genome (Tierney et al. 2019; Almeida et al. 2021). Indeed, in this study we uncover 588 inhibitory interactions between 168 chemicals of concern and human gut bacteria. The majority of these chemicals were not previously reported to have any antibacterial properties. Fungicides and industrial chemicals showed the largest impact with circa 30% exhibiting anti-commensal properties. For example, conazole fungicides inhibit ergosterol synthesis, a biochemical pathway unique to fungi. Yet, we find that conazole fungicides inhibit some Firmicutes - prominent members of the human gut microbiota.

The *in vitro* approach used in this study enabled the scale and the controlled conditions required for uncovering mechanistic relationships. Yet, many factors remain unknown to extrapolate our findings to *in vivo* conditions. One open question is exposure concentration in the colon, which can vary by geographical location, diet, occupation, bioaccumulation by the bacteria and human tissues, amongst other factors. There is currently no systematic data available on xenobiotic concentrations in the human GI tract; surprisingly also for many therapeutic drugs. Our data and the associations revealed based on toxicokinetic modelling warrant conducting such studies in the future. It will also be important to build tools and methods - both laboratory and epidemiological - towards assessing *in vivo* effects in terms of microbiome composition and function. Towards this, we used synthetic communities and these could be further expanded to include *in vitro* models like gut-on-chip (Moossavi et al. 2022). There are several insights that could be adopted from drug-microbiome interaction studies (Scher et al. 2020); particularly the power of mechanistic insights to navigate the complex *in vivo* conditions.

Towards understanding the mechanistic basis of the chemical-bacteria interactions, we used transposon insertion sequencing to determine the fitness of *P. merdae* mutants in the presence of ten xenobiotics. This revealed several efflux pumps as genetic elements linked to negative fitness which mechanistically supports the idea of a general resistant phenotype and that xenobiotics could lead to collateral resistance to antibiotics via efflux enhancement (Murray et al. 2024). Interestingly, gene hits for selection under many xenobiotics are enriched in catabolic and biosynthetic genes. This enrichment is reminiscent of the prevalence of metabolic hits observed for non-antibiotic pharmaceuticals in *E. coli* (Noto Guillen et al. 2024). The metabolism of *P. merdae* thus seems to have a major role modulating the fitness under xenobiotics. Notably, loss of genes for producing accessory metabolites with known impact on human health provided growth advantage under several xenobiotics. This raises the possibility that exposure to chemical pollutants could impact the selection landscape in the gut leading to metabolic alterations affecting the host. As these metabolic genes are well conserved and widespread in several gut bacteria these mechanisms of selection could also be present across other taxa.

Similar to the off-purpose and off-target effects of chemical pollutants, our previous work on pharmaceutical drugs (Maier et al. 2018) had revealed anti-gut bacterial effects of human targeted drugs. Taken together, these studies question the categorical classification used to label antibiotics, human-targeted drugs, industrial chemicals, and pesticides. Supporting this, we observed a linear relation between sensitivity of commensal gut bacteria to chemical pollutants and that to human-targeted drugs. The activity of drugs, for example in the gut microbiota but possibly beyond, could therefore be affected by exposure to pesticides and industrial chemicals.

While the chemical coverage of our study is comprehensive and the largest to-date, the chemical universe is vast, with chemicals being synthesised and approved for use at a fast pace. A major challenge going forward will be how to identify potential harmful compounds using computational approaches. The data from our previous drug screen (Maier et al. 2018) has been used to train machine learning models to predict bacterial toxicity from chemical features (McCoubrey et al. 2021). However, the accuracy and generality of these models is far from what will be needed from a regulatory viewpoint. Our expanded dataset increases the coverage of the xenobiotic chemical space, providing a better platform towards improving computational predictions of toxicity. Considerable expansion of the number and diversity of compounds tested also opens opportunities to gain mechanistic insights into the off-purpose and off-target antibacterial activity of chemicals. As we show in this study, general principles of xenobiotic resistance emerge: more resistance to therapeutic drugs means more resistance to chemical contaminants as well, indicating a general ‘xenobiotic resistance/sensitivity’ phenotype. Secondly, we find that more abundant strains are more susceptible, indicating a growth vs resistance trade off. A possible explanation could be that strains featuring high *in vivo* abundance have evolved to uptake and utilise diverse nutrient compounds from their environment, which also makes them more susceptible to uptake of non-nutrient compounds.

Overall, our data shows that agricultural and industrial chemicals unexpectedly affect the growth and metabolic physiology of commensal gut bacteria. The inhibition of specific bacterial species has the potential to alter the composition of the gut microbiome and therefore its vital function in human health and disease. We furthermore show that specific mutants are selected for under xenobiotic exposure, raising the possibility that such compounds affect the evolution of commensal bacteria with potential knock-on effects on antimicrobial resistance and metabolic output of the bacteria. Our study thus brings forward the gut microbiome as a site of action of chemical pollutants with implications for chemical risk assessment and understanding microbiome dynamics due to widespread chemical exposure.

## Materials and Methods

### Chemical library

An arrayed library of pesticides, pesticide metabolites and pesticide-related compounds was obtained from the Chemical Biology Core Facility at EMBL (Heidelberg, Germany). Original compound stocks were ordered from the collection of PestAnal standards by Merck/Sigma-Aldrich, with catalogue numbers and CAS identifiers listed in **Supplementary Data 1**. An additional plate containing industrial chemicals and mycotoxins was prepared in-house. Compounds were dissolved in DMSO at a concentration of 10 mM. The combined library was arranged on 12 96-well plates, which included 8 DMSO controls interspersed within the plate. One unique well position of each plate contained a dye (Trypan blue) serving as a footprint to detect and prevent plate mix ups and rotation. A working stock concentrated at 2mM in DMSO was prepared and aliquoted. Libraries were stored at −80°C.

Extensive meta-data for the library was compiled from multiple sources. The majority of PubChem CIDs were retrieved automatically by searching for the compound name using the REST PUG interface. For compounds where the search failed, IDs were added manually to the database. PubChem metadata, such as molecular formula and elemental composition, molar mass, polar area, hydrogen bond acceptors and donors, as well as InChI and SMILES structure representations for all compounds, were then retrieved based on CIDs. The number of PubMed hits for each compound in the library was retrieved in April 2022 using the NCBI E-Utilities API. We also semi-automatically matched compound names to the Pesticide Compendium of the BCPC which allowed us to map compounds to their hierarchical target classification system. For compounds that did not map to entries in the BCPC database, we manually curated metadata for pesticide metabolites and non-pesticide uses. FDA and EFSA pesticide monitoring program reports from 2018 were used to annotate the library for pesticide detection in common foods. MRL (maximum residue levels) exceedances were retrieved from the pesticide residue monitoring reports from the FDA and EFSA (2018).

### Bacterial strains and growth conditions

Details of bacterial strains, including their accession numbers in common culture collections and alternative species names, are listed in **Supplementary Data 2**. All bacteria were grown in anaerobic conditions in a polyvinyl chamber (Coy Laboratory Products) filled with 2 % hydrogen and 12 % carbon dioxide in nitrogen. The chamber was equipped with a palladium catalyst system for oxygen removal, a dehumidifier and a hydrogen sulfide removal column. Bacteria were grown at 37 °C in modified Gifu anaerobic medium (mGAM, HyServe, Germany, produced by Nissui Pharmaceuticals), prepared according to the instructions from the manufacturer and sterilised by autoclaving. Bacterial strains were chosen based on their abundance and prevalence in the healthy human gut (Dai et al. 2022).

### High-throughput screening and MIC determination

#### Growth screening

Assay plates were prepared by first diluting the frozen pesticide library aliquots (2 mM in DMSO) in mGAM within a master deep-well plate from which 50μl were aliquoted into clear round bottom polystyrene un-treated microplates (Corning 3795). Assay plates were placed in the anaerobic chamber overnight to become anoxic. All liquid handling was performed using a Biomek i7 Automated Workstation (Beckman Coulter).

Each strain was measured in biological triplicates from independent cultures inoculated on separate days. Bacteria were grown for one or two days (depending on growth rate) in 10ml of media, which were inoculated directly from frozen glycerol stocks. Cultures were then diluted 100-fold and incubated again for the same amount of time. Optical densities (OD, 600nm) were determined and a dilution with OD 0.1 was prepared. 50μl of this dilution were then added to each well of the previously prepared assay plates, resulting in a starting OD of 0.05 and a compound concentration of 20μM (1 % DMSO). Assay plates were sealed with a gas-permeable membrane (Breath-Easy, Merck, Cat# Z380059), which was additionally pierced with a syringe to prevent the build-up of gas, which otherwise occurs for some species. Plates were stacked without lids and the OD600 was recorded every hour for 24 hours using a stacker (Biostack 4, Agilent BioTek) and plate reader (Epoch 2, Agilent BioTek).

#### Statistical analysis of growth

Data analysis was performed in R version R 4.2.2 and RStudio Version 1.3.1093. First, for each growth curve the minimum OD value was set to 0. Then the raw AUC was calculated for each well using ‘bayestestR’ package and area_under_curve function. Further processing of growth curves was done by plate. For the pesticide library, AUC values were normalised by median AUC of all control wells on the respective plate. For the industrial chemicals and mycotoxin library, AUCs were normalised by median row and column values (to correct for edge effects). Then, for both screens, a z-score was calculated (raw-AUC - control median)/control sd. Each z-score was converted to its respective p-value and the p-value was FDR corrected for the number of compounds tested (n = 1076). Significance analysis was performed separately for each replicate. A replicate was considered to be significant for growth inhibition if adjusted p-value < 0.05 and > 20 % reduction in normalised AUC. A compound was considered to significantly inhibit a specific strain, if at least 2 out of 3 replicates were significant.

#### Minimum inhibitory concentration (MIC) determination

The results from the MIC screen were prepared and analysed in the same way as the high-throughput screen. The MIC was defined as the lowest concentration for which the normalised AUC dropped below 0.1.

### Comparison of compound features

The SMILES for all compounds in PubChem were downloaded, from which 250,000 were selected at random. Extended-Connectivity Fingerprints (ECFPs) were generated from SMILES for all library compounds, previously screened pharmaceutical drugs (Eskola et al. 2020; Periwal et al. 2022) and PubChem compounds, using ChemPlot (Cihan Sorkun et al. 2022). The ECFPs were then embedded in a 2 dimensional representation using Uniform Manifold Approximation and Projection (UMAP) (McInnes, Healy, and Melville 2018) with the number of neighbours and minimum distance set to 80 and 0.35, respectively. The Jaccard/Tanimoto index between compounds was calculated using the cdist function from scipy (Virtanen et al. 2020). The distributions for the maximum Tanimoto similarity index for each PubChem compound for the pharmaceutical drugs and the combined pharmaceutical drugs and pollutants libraries were compared using a two-sided KS-test.

### Compound testing in synthetic community Com20

Com20 underwent inoculation from a frozen ready-to-use stock in 5ml mGAM under anaerobic conditions for two overnight passages (diluted 1:100). This cryo stock contained each member of Com20 in equal ratios, frozen in 10% Glycerol containing Palladium black (Sigma-Aldrich, cat. No. 520810). OD578 readings were taken and Com20 was diluted to a total OD578 of 0.0125. Then, 400 µL of the diluted community was dispensed into wells of 96-deep-well plates containing xenobiotic compounds, achieving a starting OD578 of 0.01 (with a total volume of 500 µL). After sealing the deep-well plates with an AeraSeal breathable membrane (Sigma-Aldrich, cat. No. A9224), they were anaerobically incubated at 37 °C for 24 hours. Pellets from 300 µL of the cultures were frozen for subsequent 16S rRNA gene analysis and OD578 was measured as a proxy for total biomass.

DNA extraction was carried out from 300 μL culture pellets using the DNeasy UltraClean 96 Microbial Kit (Qiagen 10196-4). Subsequent library preparation and sequencing were conducted at the NGS Competence Center NCCT (Tübingen, Germany). Genomic DNA was quantified using the Qubit dsDNA BR/HS Assay Kit (Thermo Fisher) and adjusted to 100 ng input for library preparation. The initial step PCR was performed in 25 µL reactions containing KAPA HiFi HotStart ReadyMix (Roche), 515F51 and 806R52 primers (targeting a ∼350-bp fragment of the 16S V4 region), and template DNA (PCR program: 95 °C for 3 min, 28 cycles of (98 °C for 20 s, 55 °C for 15 s, 72 °C for 15 s), 72 °C for 5 min). Following PCR, the products were purified using 28 µL of AMPure XP beads and eluted in 50 µL of 10 mM Tris-HCl. Indexing was then performed in a second step PCR using KAPA HiFi HotStart ReadyMix (Roche), index primer mix (IDT for Illumina DNA/RNA UD Indexes, Tagmentation), and the purified initial PCR product as template (PCR program: 95 °C for 3 min, 8 cycles of (95 °C for 30 s, 55 °C for 30 s, 72 °C for 30 s), 72 °C for 5 min). Following another round of bead purification (20 µL of AMPure XP beads, eluted in 30 µL of 10 mM Tris-HCl), the libraries were evaluated for correct fragment length on an E-Base device using E-Gel 96 Gels with 2% mSYBR Safe DNA Gel Stain (Fisher Scientific), quantified with a QuantiFluor dsDNA System (Promega), and then pooled equimolarly. The pooled libraries were subsequently sequenced on an Illumina MiSeq device with a v2 sequencing kit (input molarity 10 pM, 20% PhiX spike-in, 2×250 bp read lengths).

We employed the DADA2 v. 1.21.053 package within R (version 4.2.0), adhering to its standard operating procedure outlined at https://benjjneb.github.io/dada2/bigdata.html. In summary, after scrutinising the quality profiles of the raw sequences, we applied trimming and filtering to the paired-end reads using specific parameters: trimLeft: 23, 24; truncLen: 240, 200; maxEE: 2, 2; truncQ: 11. Subsequently, the filtered forward and reverse reads underwent separate dereplication and were utilised for the inference of amplicon sequence variants (ASVs) using default settings, followed by merging the reads on a per-sample basis. Further, we filtered the merged reads to retain only those falling within a length range of 250 to 256 bp and conducted chimera removal. Taxonomic assignment was carried out in two stages. Initially, the final set of ASVs was classified up to the genus level utilising a curated DADA2-formatted database derived from the genome taxonomy database (GTDB) release R06-RS20253, accessible at https://scilifelab.figshare.com/articles/dataset/SBDI_Sativa_curated_16S_GTDB_database/14869077. Subsequently, ASVs belonging to genera anticipated within COM20 were further classified at the species level employing a modified version of the aforementioned database, comprising solely full-length 16S rRNA sequences of the 20 synthetic community members. Each ASV’s sequence was aligned against this database using the R package DECIPHER v. 2.24.054; classification as a specific species was assigned if the sequence similarity exceeded 98% with the closest member in the database. The abundance of each taxon within COM20 was determined by aggregating reads at the species level.

### Toxicokinetic modelling

Calculation of Estimated Administered Doses (EAD): To explore the effects of chemical pollutants on gut microbiota and extrapolate *in vitro* bioactivity data to potential human health risks, we focused on 168 active chemicals. *In vitro* concentration data from growth screening assay were used to compute EADs via the IVIVE tool from the Integrated Chemical Environment (ICE; https://ice.ntp.niehs.nih.gov/), leveraging the EPA’s httk R package (Pearce et al. 2017) and in-house code from NICEATM (Bell et al. 2020). IVIVE calculations involved determining concentrations in the gut corresponding to bioactivity levels (MICs from growth screening assays). Essential parameters included the toxicokinetic model type, chemical-specific parameters (logP, molecular weight, pKa, intrinsic clearance, fraction unbound, and tissue partition coefficient), dosing scenarios, and exposure routes. These parameters were computed using OPERA (Mansouri et al. 2018) for 152 of the 168 chemicals, and were empirically available for the remainder. The PBTK Model utilises multiple compartments for various organs and tissues, employing perfusion rate-limited kinetics to simulate elimination primarily via hepatic and renal routes, here under an oral exposure scenario for 30 days with 24-hour intervals. The model includes compartments for the gut, liver, and remaining body, simulating dynamic tissue and plasma concentrations for both oral and intravenous exposures (Chang et al. 2022).

Curation of *in vivo* POD data: The U.S. EPA ToxValDB served as the primary source for condensed *in vivo* POD data, encompassing human health references and animal toxicity values from various studies, accessible via the CompTox Chemicals Dashboard (https://comptox.epa.gov/dashboard/). Filters applied to the data available for 126 chemicals included exposures in mg/kg-bw or mg/kg-bw/day units, or convertible units, species-specific conversion factors for rats, mice, dogs, and rabbits, and study types “sub-chronic”, “subacute”, and “subchronic.” Only LOEL, LOAEL, NOEL, and NOAEL POD types were considered.

### TnBarSeq assay and analysis

The dense transposon mutant library of *P. merdae* ATCC 43184 (manuscript in preparation) was used for Barcoded Transposon Sequencing (TnBarSeq) mapped to the annotated genome GenBank: CP102286.1. Two vials of the pooled stock Tn library were used to inoculate mGAM and grown until mid-exponential phase. This culture was used to inculcate at OD_600_=0.02 1 mL cultures in a 2 mL deep well plate, with duplicates for each xenobiotic and triplicates for the respective control media with DMSO 1 %, DMSO 0.2 % or plain media (xenobiotic concentrations at **Supplementary Data 6**). The growth of the library was tracked by monitoring hourly a 100 uL aliquot of the library (as described above). The Tn library cultures were grown to early stationary phase (16 h for all samples except closantel 2.5 μM grown to 48 h). Genomic DNA was extracted with MagMAX Microbiome Ultra Nucleic Acid Isolation Kit. The transposon barcodes were amplified by PCR with NEB Q5 Hot Start HF X2, using indexed primers, and sequenced with Illumina NextSeq 75 nt SE. Barcodes were counted with 2FAST2Q (Bravo, Typas, and Veening 2022; de Bakker et al. 2022) and changes in mutant strain abundance were analysed with TRANSIT (DeJesus et al. 2015) against their respective control (**Supplementary Data 6**). KEGG terms were assigned with EggNOG Mapper (Cantalapiedra et al. 2021) and KEGG term enrichment was performed with a Fisher’s Exact Test in R. Protein homology was mapped with clinker (Gilchrist and Chooi 2021), membrane protein families were annotated with TransAAP at TransportDB and Conserved Domains Database at NCBI (Elbourne et al. 2023) and additional relevant homologs were identified by PaperBLAST (Price and Arkin 2017).

## Data Availability

The following supplementary files are associated with this manuscript: **Supplementary Data 1** - Chemical compound library meta-data **Supplementary Data 2** - Bacterial strain details **Supplementary Data 3** - Growth screen results **Supplementary Data 4** - Community experiment results **Supplementary Data 5** - Toxicokinetic modelling **Supplementary Data 6** - TnBarSeq results All code for the generation and comparison of ECFPs is available in a github repository: https://github.com/MRCToxBioinformatics/Compound_bacteria_screen_chemical_spacev0.1.0. The TnBarSeq data for this study has been deposited in the European Nucleotide Archive (ENA) at EMBL-EBI under accession number PRJEB76605.

## Acknowledgements

We would like to thank Carlos G P Voogdt and Athanasios Typas for providing materials and guidance for TnBarSeq library construction. We thank the Chemical Biology Core Facility at EMBL (Heidelberg, Germany) for assembling the pesticide library. We thank EMBL Genecore (Heidelberg, Germany) for their support with sequencing of the TnBarSeq samples. This project has received funding from the European Research Council (ERC) under the European Union’s Horizon 2020 research and innovation programme (grant no. 866028) and from the UK Medical Research Council (project no. MC_UU_00025/11). L.M. was supported by the DFG (EXC2124, MA 8164/1-2) and the ERC (gutMAP, 101076967). This research was supported in part by the Intramural Research Program of the NIH, National Institute of Environmental Health Sciences (NIEHS).

