## Supplementary Figures 1-4 for "Off-purpose activity of industrial and agricultural chemicals against human gut bacteria"

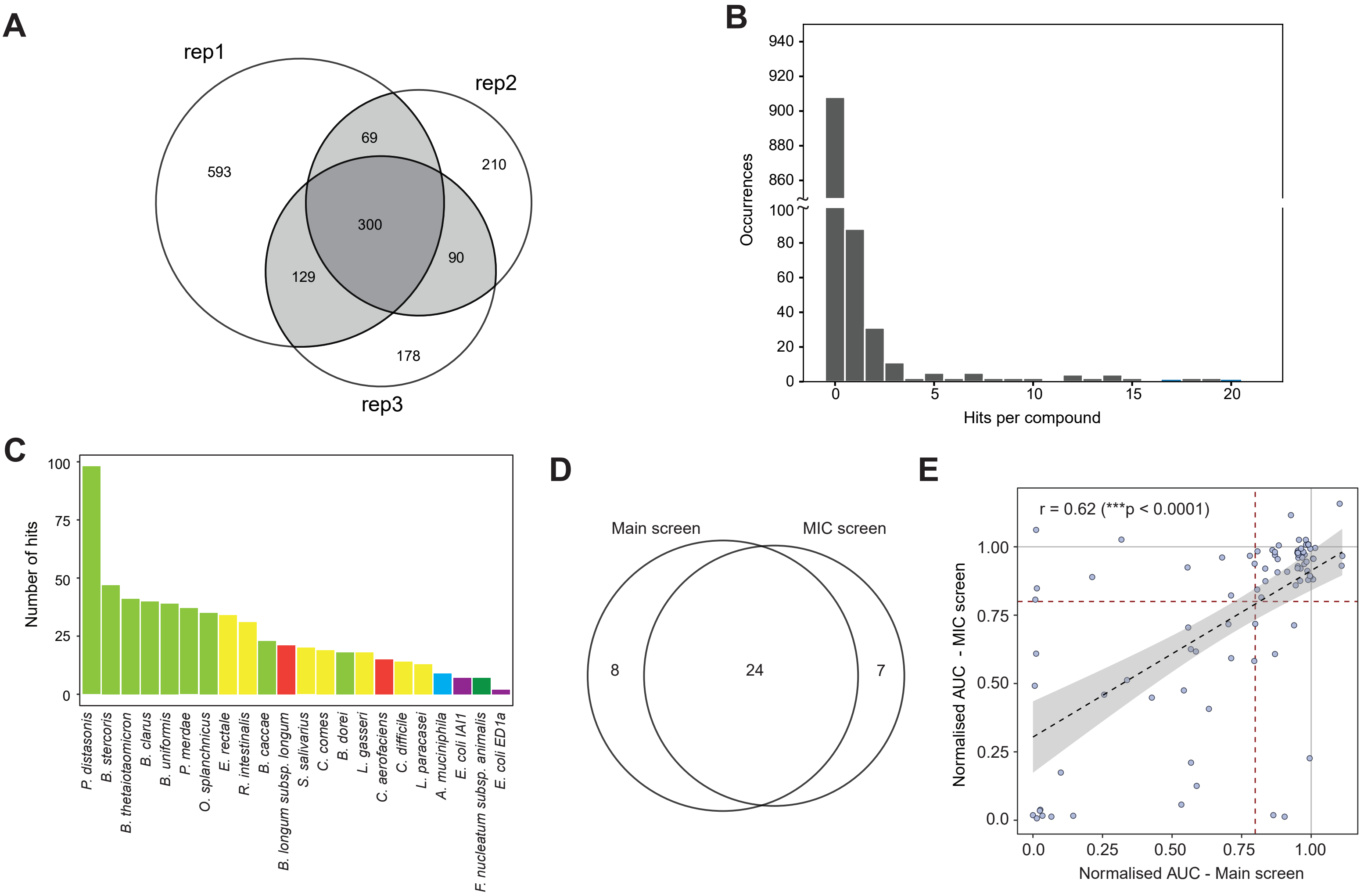


**Supplementary Figure 1:**

A) Venn diagram illustrating overlaps of significant interactions between n=3 biological replicates. A compound was considered an overall hit if two out of three biological replicates were statistically significant ($p_{adj}<0.05$) and the area under the curve (AUC) was at least 20% lower than for the DMSO control.

B) Histogram illustrating number of hits per compound.

C) Number of hit compounds per species.

D) Results of the screen were validated in a separate screen with independently obtained and prepared compounds. The Venn diagram illustrates the overlap of hits between the two screens.

E) Normalised AUC values in main screen and validation experiment. Pearson correlation is shown in the graph.

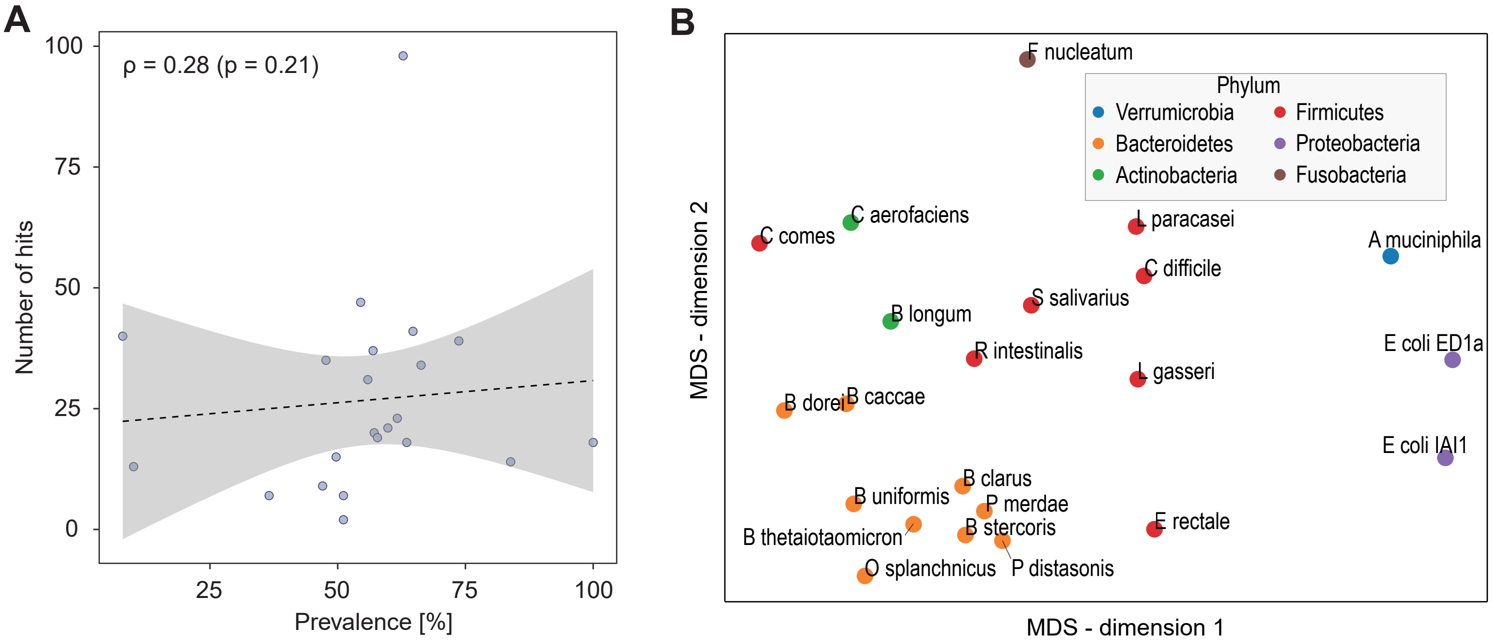


**Supplementary Figure 2:**

A) Correlation analysis between prevalence of bacterial species across human microbiomes and number of hits in growth inhibition screen.

B) Multi-dimensional scaling of bacterial susceptibility profiles indicates that genetically more closely related species have more similar profiles, indicating a genetic basis for susceptibility. The distance matrix was computed using Jaccard index and multi-dimensional scaling was performed in scikit-learn with default settings.

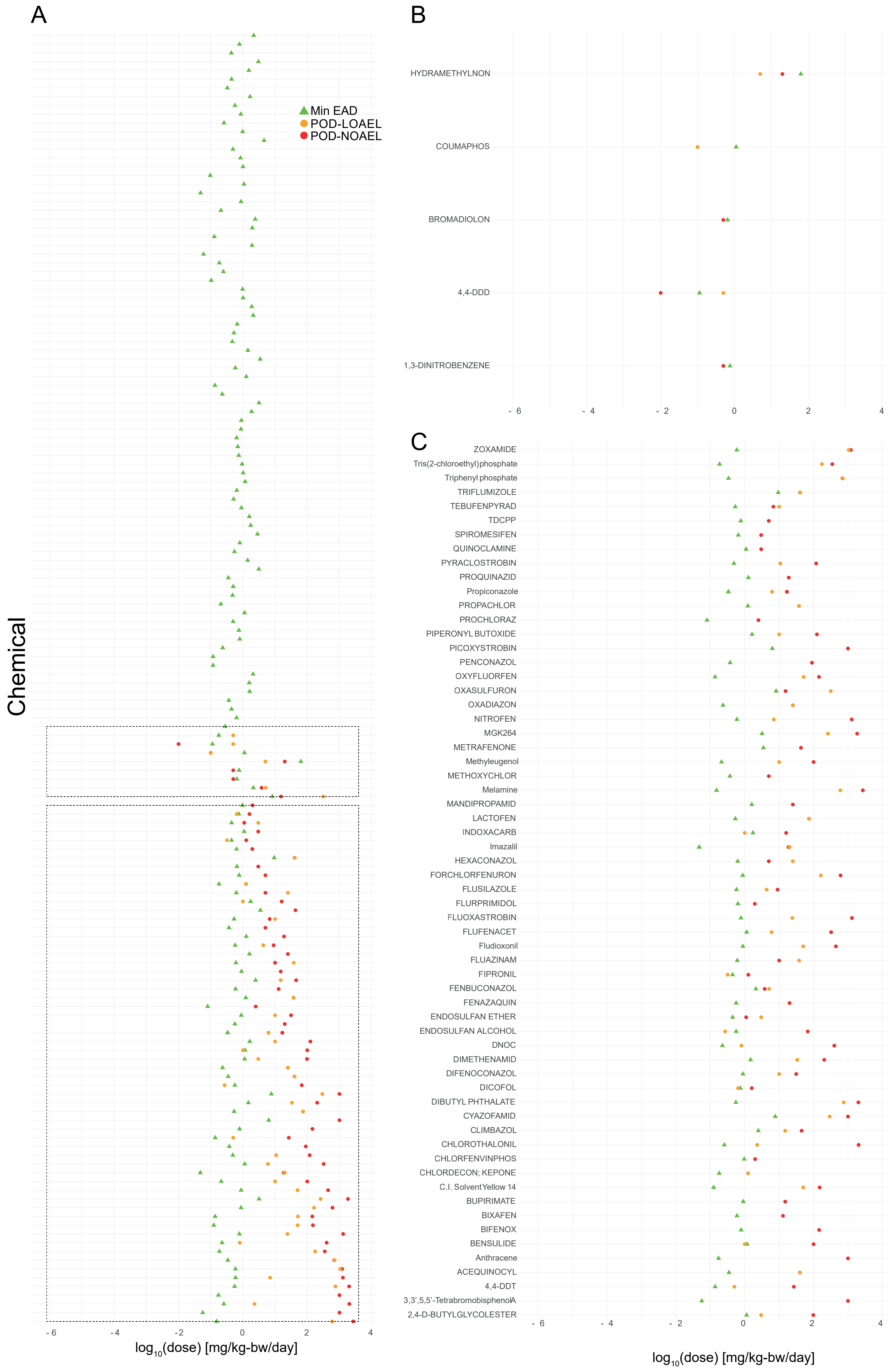


**Supplementary Figure 3:**
Toxicokinetic modelling and *in vitro* to *in vivo* extrapolation (IVIVE). Analysis of minimum equivalent administered oral doses in humans EAD (green triangle), Point of departure - lowest adverse effect level (POD-LOAEL,orange circle) and Point of departure - no adverse effect level (POD-NOAEL,orange circle), for 149 Chemicals. Boxes highlight chemicals with negative and positive POD ratios, enlarged in panel B and C respectively.

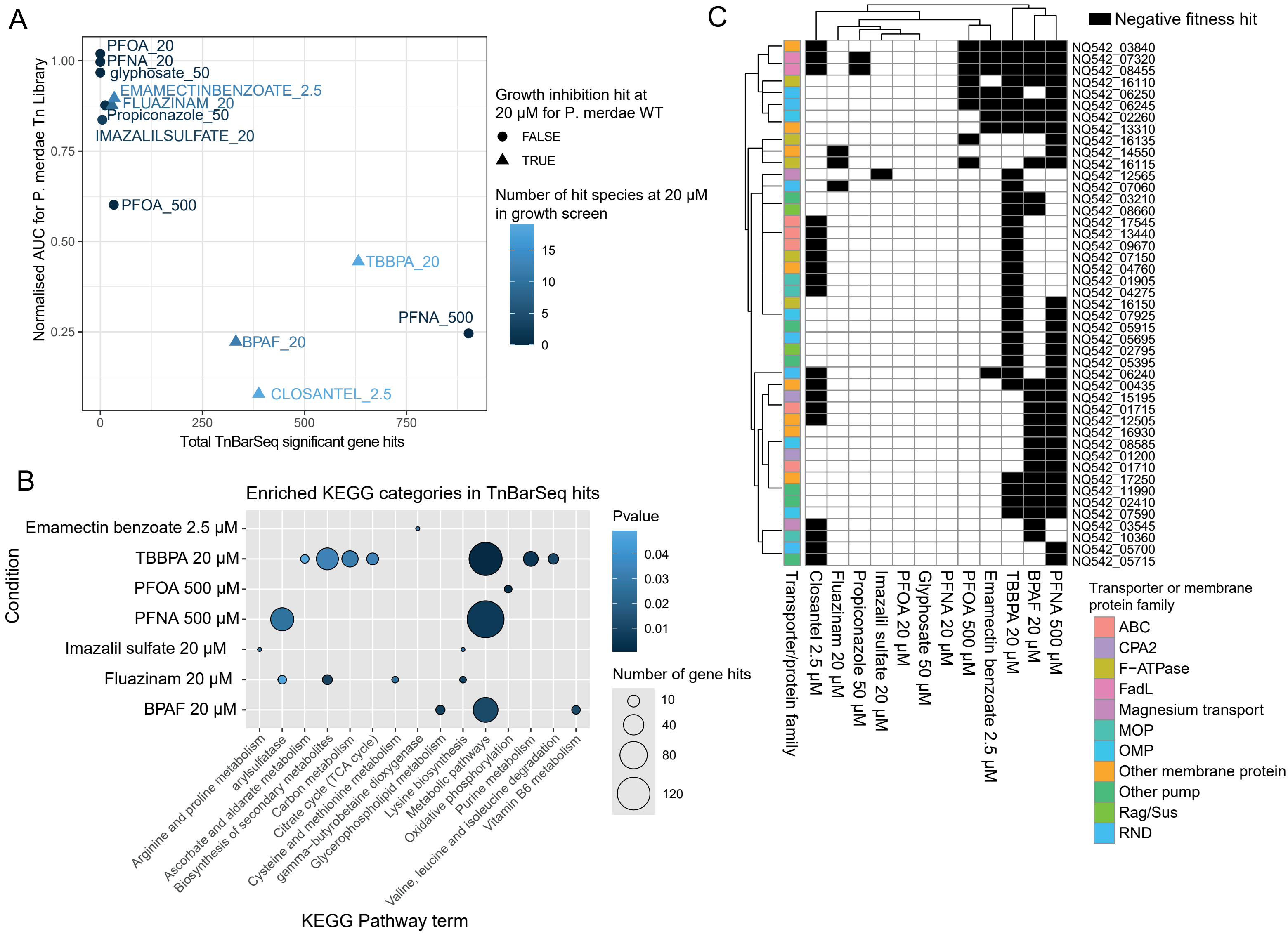

**Supplementary Figure 4:**

A) Normalised area under the Tn mutant library growth curve (AUC) for the compounds tested vs total Tn-Barseq-hits. The number of bacterial species that were a hit in the growth inhibition screen for the compound at 20 μM is indicated in colour scale. The point shape indicated if the compound at 20 μM was a hit for WT *P. merdae* in the previous growth screen.

B) Enriched KEGG KO categories in all TnBarSeq hits show enrichment of several metabolic pathways. Number of gene hits are indicated by circle size.

C) Putative membrane transporters and other membrane proteins that are negative fitness hit in more than one xenobiotic. Protein families are indicated in colours. ABC= Adenosine Triphosphate-Binding Cassette. CPA2= Monovalent cation:proton antiporter 2. RND= Resistance Nodulation Division. MOP= Multidrug/oligosaccharidyl-lipid/polysaccharide. omp= ompA-H family.
